# Inferring metabolite interactomes via molecular structure informed Bayesian graphical model selection with an application to coronary artery disease

**DOI:** 10.1101/386409

**Authors:** Patrick J. Trainor, Joshua M. Mitchell, Samantha M. Carlisle, Hunter N.B. Moseley, Andrew P. DeFilippis, Shesh N. Rai

## Abstract

**Introduction:** While the generation of reference genomes facilitates the elucidation of gene-phenome associations, reference models of the metabolome that are specific to organism, sample type (e.g. plasma, serum, urine, cell-culture), and state (including disease), remain uncommon. In studying heart disease in humans, a reference model describing the relationships between metabolites in plasma has not been determined but would have great utility as a reference for comparing acute disease states such as myocardial infarction.

**Materials and Methods:** We present a methodology for deriving probabilistic models that describe the partial correlation structure of metabolite distributions (“interactomes”) from metabolomics data. As determining partial correlation structures requires estimating *p*(p-1)/2* parameters for *p* metabolites, the dimension of the search space for parameter values is immense. Consequently, we have developed a Bayesian methodology for the penalized estimation of model parameters in which the magnitude of penalization is drawn from probability distributions with hyperparameters linked to molecular structure similarity. In our work, structural similarity was determined as the Tanimoto coefficient of algorithmically-generated “atom colors” that capture the local structure around each atom within each structure. A Gibbs sampler (a Markov chain Monte Carlo technique) was implemented for simulating the posterior distribution of model parameters. We have made software for implementing this methodology publicly available via the R package *BayesianGLasso*.

**Results / Conclusions:** First, we demonstrate robust performance of our methodology (sensitivity, specificity, and measures of accuracy) for recovering the true underlying partial correlation structure over simulated datasets (with simulated metabolite abundances and simulated known structural similarity). We then present an interactome model for stable heart disease inferred from non-targeted mass spectrometry data via this methodology. Inspection of the local graph topology about cholate reveals probabilistic interactions with other primary bile acids, secondary bile acids, and many steroid hormones sharing the same precursors.

## 1. Introduction

Untargeted profiling of the metabolome of an organism provides a view into the small molecule determinants of phenotype. While the genome of an organism may be conceptualized as a blueprint for the composition and organization of an organism that is largely immutable (barring epigenetic modifications, DNA damage, and genetic mutations) (Gao, Jia, Zhang, Breitling, & Brenner, 2015; Keating & El-Osta, 2015; Martincorena & Campbell, 2015), the metabolome of an organism is dynamic and variable (Dallmann, Viola, Tarokh, Cajochen, & Brown, 2012; Krycer et al., 2017). Sources of variation within the metabolome of a single organism include tissue-, cell-, and organelle-specific localization of metabolic processes (Shlomi, Cabili, Herrgård, Palsson, & Ruppin, 2008; Voet, Voet, & Pratt, 2013); environmental exposures (Southam et al., 2014); and host-microbe interactions and metabolite exchange (Moriya, Satomi, Murata, Sawada, & Kobayashi, 2017). While the generation of reference human genomes has facilitated the interrogation of gene-phenome associations (including human disease associations), the intrinsic variability and dynamic nature of the metabolome of an organism likely precludes the generation of such a reference model. While a single reference model of the metabolome of an organism may not be sensible, significant efforts such as the HUSERMET project (Dunn et al., 2014) have been undertaken to quantify the repertoire of metabolites in specific biofluids for examining metabolite-metabolite and metabolite-phenotype associations. In order to make systems-level comparisons of the differences in the metabolome across phenotypes, models of the conditional relationships between metabolites are necessary. By the determination of sample media and analytical platform-specific probabilistic interaction models, henceforth called “interactomes”, systems-level comparisons of phenotypes that can be made. A specific use case for such an interactome model is the generation of a plasma interactome for stable heart disease.

Heart disease is the most prevalent cause of death globally (Benjamin et al., 2017). As a disease, heart disease does not represent a uniform condition, but rather a collection of diseases of varying etiologies (Kasper, 2015). Of particular interest in the study of coronary artery disease (CAD) is the elucidation of the precipitants of acute disease events such as myocardial infarction (Arbab-Zadeh & Fuster, 2015) or unstable angina, metabolic pathways associated with disease phenotypes (Y. Fan et al., 2016), and determining the metabolic consequences of acute events (Trainor et al., 2017). To date, an interactome describing the conditional relationships between blood plasma metabolites in humans with heart disease does not exist. If such a reference model were determined, it would facilitate making systems-level inferences regarding metabolic perturbations that accompany acute disease events such as unstable angina or acute myocardial infarction (MI).

While correlation networks have been used to describe the relationships between metabolites in many metabolomics experiments [see for example (Kotze et al., 2013; Madhu et al., 2015; Suarez-Diez et al., 2017; L. Wang et al., 2015)], this approach is limited as the topology learned represents only the pairwise marginal associations between metabolites. Determining a conditional relationship between two metabolites allows for inference regarding how the abundance of a specific metabolite influences the abundance of another metabolite after conditioning on the abundance of other intermediates. In order to model such conditional probabilistic dependencies between metabolite abundances, a Gaussian Graphical Model (GGM) approach may be employed as in the present work. GGMs provide a suitable framework for representing the joint probability distribution of metabolites that are detected in metabolomics experiments and for representing the probabilistic interactions between metabolites and have been employed for such a task previously (Krumsiek, Suhre, Illig, Adamski, & Theis, 2011; Shin et al., 2014).

A significant challenge in evaluating the relationships between metabolites in an untargeted metabolomics experiment is that the dimension of metabolites may be greater than the number of samples. Even given a relatively high ratio of samples to metabolites detected, in the evaluation of pairwise conditional relationships between metabolites, the number of parameters to be estimated can be prohibitive. For example, if *p* = 500 metabolites are detected, an evaluation of all pairwise conditional relationships would require the simultaneous estimation of 124,750 parameters. The use of regularization is a well-established approach for guaranteeing the existence of Gaussian Graphical Model parameters, amenable to the case that the sample size *n* is less than *p* (Banerjee, El Ghaoui, & d'Aspremont, 2008; J. Fan, Feng, & Wu, 2009; Friedman, Hastie, & Tibshirani, 2007; Meinshausen & Bühlmann, 2006; Yuan & Lin, 2007).

Penalized estimation of GGM parameters provides a natural mechanism for integrating *a priori* knowledge regarding the molecular structure of metabolites with experimental metabolomics data. The integration of empirical data and scientific knowledge regarding metabolism is common in metabolomics studies. Typically, univariate and/or multivariate analyses first identify sets of metabolite features for which evidence of differences between experimental conditions or phenotypes are observed. After identifying interesting metabolite features, these sets are tested for enrichment of specific metabolic pathways or biological processes greater than that expected by chance (Xia & Wishart, 2010). A promising alternative to pathway analyses discussed in (Dinesh Kumar g169
Barupal & Fiehn, 2017) is to use structural similarity and chemical ontology as *a priori* knowledge to generate study-specific metabolite sets for contextualizing empirical results. The current work is of a similar paradigm and predicated on the assumption that the individual biochemical reactions that result in statistical dependence between metabolic intermediates also generate statistical dependence in structural similarity between the same intermediates. In our application, structural similarity is determined by an approach that considers overlap in shared local structure between metabolites. Rather than considering fixed sets of metabolites such as pathways, sets, or modules and subsequently quantifying enrichment of these sets in empirical results, we consider *a priori* knowledge of the relationships between metabolites as probabilistic statements about the relatedness of compounds. Thus, the *a priori* scientific knowledge is used to generate prior probability distributions that influence GGM model selection, so that posterior inference probabilistically combines empirical data and prior scientific knowledge to yield an updated model of the probabilistic interactions between metabolites. In the present work, we introduce a methodology for using molecular structure similarity to generate prior distributions that control the degree of penalization in parameter estimation for learning a GGM metabolite interactome from metabolomics data.

We first evaluated the methodology using simulation studies. For these simulation studies, autoregressive processes were simulated for representing linear biological processes in which the correlation between simulated metabolites decreased in tandem with decreasing structural similarity. We evaluated the sensitivity, specificity, AUC, and *F_1_* measure of the proposed method in recovering the true pairwise conditional correlations structures that were specified in advance. Finally, we applied our methodology to a human plasma dataset, specifically for the development of a reference model for stable heart disease.

## 2. Methods

### 2.1. Gaussian Graphical Models (GGM)

We consider Markov Random Fields (MRFs) which are graphical models in which random variables *X_i_* ∈ *V, i* = 1, 2, …, *p* are represented as vertices and edges in the edge set *E* ⊆ *V* × *V* represent probabilistic interactions. Gaussian Graphical Models (GGMs) represent a special class of MRFs in which the underlying joint probability distribution represented by the graph is assumed to be multivariate Gaussian (Koller & Friedman, 2009). In addition to the joint distribution being multivariate Gaussian, the marginal distribution for each *X_i_* is Gaussian, as are the conditional distributions for *X_i_*|*X_j_*. Given a multivariate Gaussian distribution 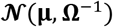 where **μ** is a vector of means and **Ω** is the inverse of the covariance matrix **Σ** (i.e. a concentration matrix), the entries *ω_ij_* of the concentration matrix are of particular importance as *ω_ij_* = 0 implies that *X_i_* and *X_j_* are conditionally independent and with respect to the graph topology, there does not exist an edge between *X_i_* and *X_j_*. Further from the entries of Ω, the partial correlation coefficient between two random variables *X_i_* and *X_j_* can be computed as: 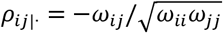

### 2.2. GGM parameter estimation

It has been shown previously that if *n* < *q* where *q* represents the maximal clique size of the GGM then a maximum likelihood estimator does not exist (Buhl, 1993). Noting the likelihood function for the concentration matrix **Ω**:

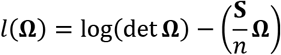

where **S** = **X**^*T*^**X** is the sum of products matrix. As the log-likelihood function is not guaranteed to be convex, regularization of this likelihood has been proposed (Banerjee et al., 2008; Friedman et al., 2007; Meinshausen & Bühlmann, 2006) as a solution for estimating **Ω**. Friedman et al. (2007) proposed a method, known as the graphical Lasso (Least Absolute Shrinkage and Selection Operator) for finding the maximum of the *L_1_*norm penalized log-likelihood:

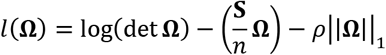

via a coordinate descent algorithm.

A Bayesian approach has been proposed for the regularized estimation of **Ω** (H. Wang, 2012) that provides a natural structure for integrating *a priori* scientific knowledge and high-throughput molecular biology data such as untargeted metabolomics data. H. Wang (2012) introduced a hierarchical Bayesian representation of the regular graphical Lasso as well as the adaptive graphical Lasso (J. Fan et al., 2009). The frequentist adaptive graphical Lasso was devised to link the magnitude of the penalty parameter to the norm of individual concentration matrix entries and proposes the following penalized likelihood for **Ω**:

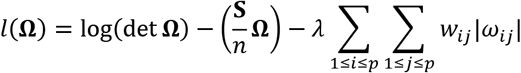

with weights 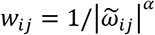 where *α* > 0 and 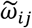 are estimates for the concentration matrix entries, such as regular graphical Lasso estimates.

As a Bayesian model:

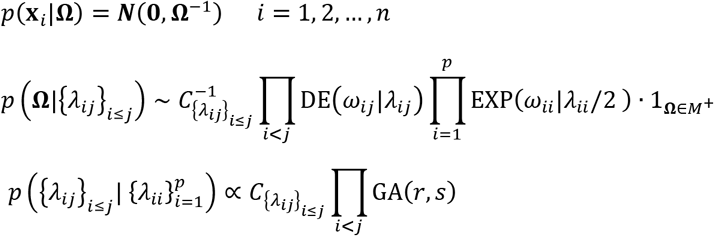

In the above model formulation for the density of **Ω** conditional on the *λ_ij_*, DE(·|*λ_ij_*) represents the double exponential, or Laplace distribution, and EXP(·|*λ_ij_*) the exponential distribution with scale parameter *λ_ij_*. The space of positive definite matrices is represented by 1_**Ω**∈*M*^+^_. Finally, *C* represents the normalizing constant so that *p*(**Ω**|{*λ_ij_*}_*i*≤*j*_) is a proper probability distribution. For the non-adaptive Bayesian graphical Lasso, *λ_ij_* = *λ* for all *i* and *j*, in other words the shrinkage parameter is not specific to each concentration matrix entry. H. Wang (2012) chooses a gamma prior for *λ_ij_*, that is *λ_ij_* ~ Gamma(*r, s*), where *r* and *s* are hyperparameters and develops a data-augmented block Gibbs sampler for sampling from the posterior distribution of **Ω**. Further, it is shown that the conditional distribution of the shrinkage parameter is then 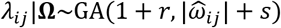. In this case, the scale hyperparameter for the shrinkage varies with the norm of the current MCMC iteration estimate 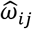 for each *ω_ij_* allowing for adaptive penalization as introduced by J. Fan et al. (2009). We propose that to incorporate prior knowledge regarding the relatedness of compounds, the scale hyperparameter can be linked to structural similarity, that is by specifying the prior distribution *λ_ij_* ~ Gamma(*r, s_ij_*) where *s_ij_* is a measure of structural similarity between compound *i* and compound *j*. The conditional expected value of each *λ_ij_* is then: 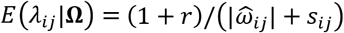.

### 2.3. Generating informative priors from molecular structure

To generate informative shrinkage priors for the adaptive Bayesian graphical Lasso, we utilized a local structure similarity metric. This metric was adapted from the previously described Chemically Aware Substructure Search (CASS) algorithm (Mitchell, Fan, Lane, & Moseley, 2014). In this adaptation, the structural similarity between any two chemical structures (*A* and *B*) was estimated using strings representing local chemical structure (referred to as the atom’s color) centered at every atom in the two structures. The color of every atom was constructed as follows. First, for every bonded atom, its element type and the order of the bond connecting it to the center atom are joined to form a component string that is added to a list of components. For example, if the center atom has a double bonded oxygen, this would contribute a ‘O2’ component to the component list. Every component represents a portion of the local bonded structure at the center atom. Second, the components strings are then sorted alphanumerically and concatenated to produce a description of the bonded structure one bond away from the center atom. Finally, to the front of this string, the element type of the center atom is then added to yield the atom’s color. Each color uniquely maps to a single locally bonded structure (e.g. the ‘CC1O1O2’ coloring represents a carbon of a carboxylate). Since the component list was first sorted alphanumerically, this color is consistent for all identical local structures regardless of how they are ordered in their representation. Each chemical structure can be represented as the list of its constituent atom’s colors and these lists of colors can be compared to determine structural similarity. To determine structural similarity between compounds using the color string representations, the Tanimoto coefficient between pairs of compounds was computed, which is defined as (Chen & Reynolds, 2002):

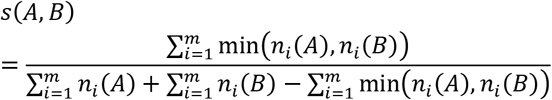

where *n_i_*(*A*) represents the count of unique colored atoms indexed by *i* = 1,2, …, *m* for molecule *A*. The Tanimoto dissimilarity is then *d*(*A,B*) = 1 – *s*(*A,B*).

After determining the Tanimoto dissimilarity between each pair of metabolites, the gamma hyperprior distribution for the shrinkage parameter *λ* can be determined by linking the gamma distribution shape to the dissimilarity, that is by setting *s_ij_* = *f*(1 – *d*(*i,j*)) where *i,j* index metabolites and *f*(*x*) is a monotonic function. The conditional distribution of the shrinkage parameter is then 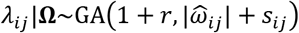. A plot of the relationship between the structural similarity of two hypothetical metabolites and the expected value of the shrinkage parameter is shown in Figure 1.

**Figure 1:**
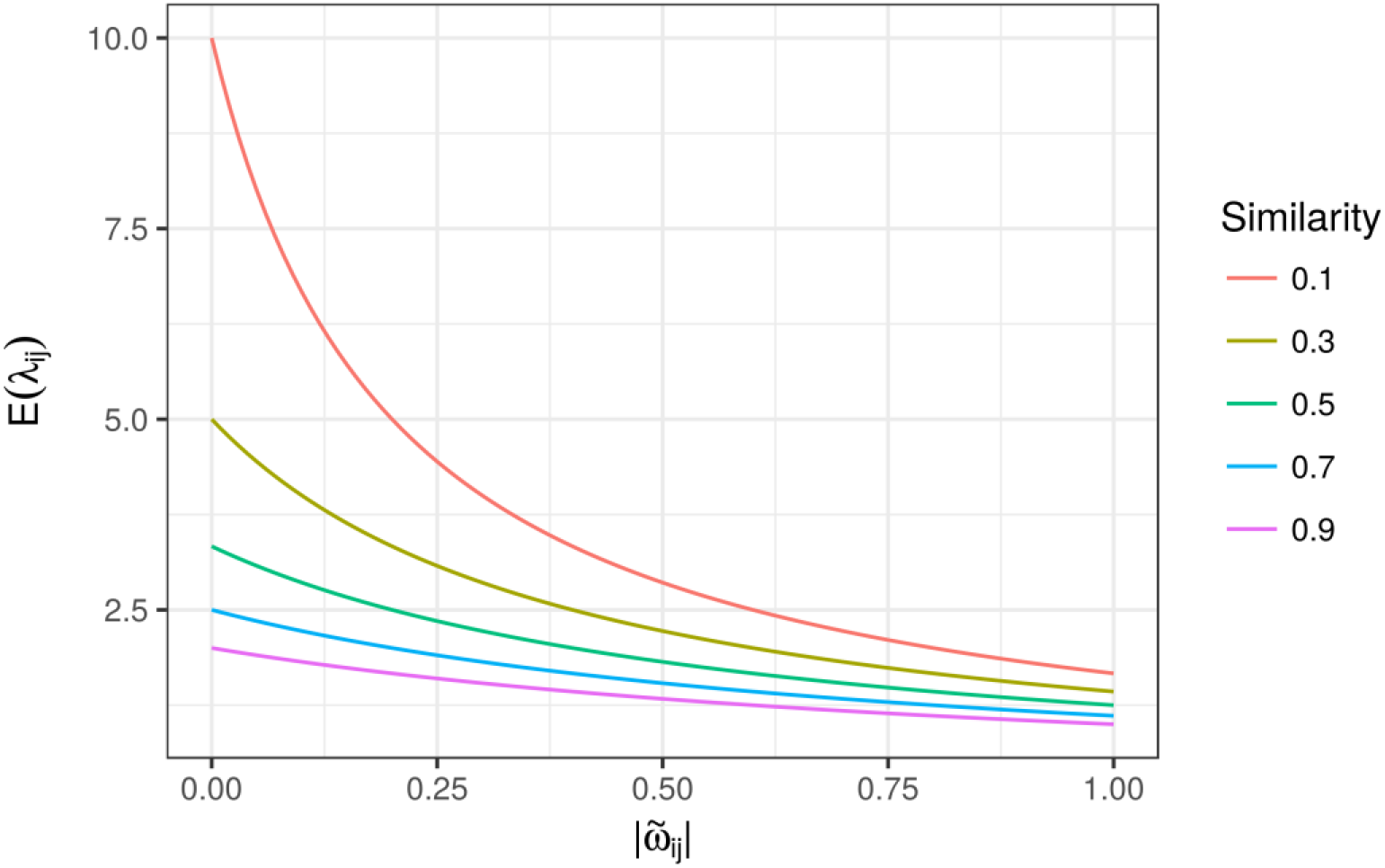
Theoretical relationship between the structural similarity of a pair of metabolites and the expected value of the shrinkage parameter AlJ-during model estimation. The horizontal axis represents a current estimate for a concentration matrix entry for a pair of metabolites. The vertical axis represents the expected value of the shrinkage parameter. Color values show the structural similarity between the pair of metabolites.

### 2.4. Posterior inference of model parameters

To determine a graphical model (or a set of models of high probability) given structural priors samples may be drawn from the posterior distribution of *p*(**Ω**|**X**), using a Gibb’s sampler similar to that introduced by Wang (2012). Previous work has shown that the exponential power family of probability distributions can be represented as a scale mixture of normal distributions with a defined mixing density (West, 1987). Using this fact and introducing the latent scale parameter *τ*, the unnormalized posterior distribution can be written as:

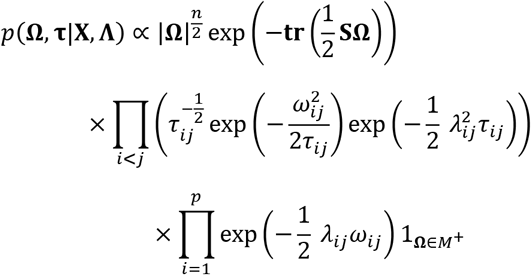

The block Gibb’s sampler cycles through column-wise partitions of **Ω**, drawing from the conditional distribution of a single column of the matrix **Ω**, conditioned on the current values of the remaining columns. We developed an R package, *BayesianGLasso*, for implementing this and other samplers for the Bayesian Graphical Lasso. The underlying sampler was written in C++ using *Rcpp* and *RcppArmadillo* to make use of the Armadillo linear algebra library. In addition to providing Gibb’s sampling methods the R package developed by our group includes classes for storing the Markov chains generated by the sampler along with relevant parameters and hyperparameters, and methods for conducting statistical inference over the simulated posterior distributions.

### 2.5. Efficacy analysis via simulation studies

To evaluate the efficacy of the proposed method, we employed simulation studies. We sought to evaluate the relative performance of the adaptive Bayesian Graphical Lasso (BGL) using informative priors versus (1) the adaptive Bayesian Graphical Lasso using non-informative priors, and (2) the Bayesian Graphical Lasso (non-adaptive). For the informative prior case, we further manipulated the degree to which the priors were accurate relative to the partial correlation structure utilized to generate the data. We evaluated the methods by simulating both simple partial correlation structures as well as more complex structures utilizing two simulation schemas. Under the first schema, a simple autoregressive (AR) process of order 1 was simulated for representing a linear biological process with decreasing structural similarity with increasing process distance. Simulated structural similarity was taken to be deterministically known, that is a structural similarity matrix was defined as: **Σ** = [*σ_i,j_*] where *σ_ij_* = *ρ*^|*i*–*j*|^. To simulate metabolite abundances, a random matrix was sampled from the multivariate normal distribution ***N***(**0,Σ**). In the “accurate” informative prior case, the shrinkage hyperprior *s_ij_* was defined as 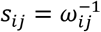, where 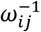 are the elements of **Ω** = **Ω**^−1^. After generating simulated datasets, the adaptive BGL (with informative and non-informative priors) as well as the non-adaptive BGL were utilized for estimating the concentration matrix and corresponding graph topology. Given the simple dependence structure in the AR(1) case, a measure of ground truth was available as the existence of edges between simulated metabolites was known *a priori*. We evaluated the sensitivity, specificity, and F1 measure of each method for detecting the presence of edges by utilizing the magnitude of the estimated concentration matrix entries: 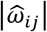. In addition, we report the area under the receiver operating characteristic curve for assessing each technique, which considers the range of possible fixed cutoff values of 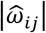 for estimating the presence or absence of edges. While each technique draws shrinkage parameters from a Gamma distribution, the shape and scale of each distribution depends both on empirical data and hyperparameters. We conducted shape and scale hyperparameter optimization separately for each technique via a grid search over simulated datasets prior to the evaluation of performance.

### 2.6. A plasma interactome for stable heart disease

In order to determine changes in the plasma metabolome associated with myocardial infarction (MI) characterized by thrombotic etiology versus non-thrombotic etiology, DeFilippis and colleagues assembled a human cohort as previously described (DeFilippis et al., 2016; DeFilippis et al., 2017; Trainor et al., 2017). Briefly, 80 human subjects presenting with suspected acute MI or stable coronary artery disease (CAD) were enrolled. Utilizing a stringent criteria based on clinical presentation, angiographic evidence, and histological evidence, MI subjects were adjudicated as thrombotic MI or non-thrombotic MI. Blood samples were collected at the time of acute presentation (presentation to the coronary artery catheterization lab prior to procedures) and at a follow-up evaluation approximately three months later. To estimate the structure of a stable heart disease plasma interactome, we used the follow-up evaluations from all available MI subjects as well as the evaluations from stable CAD subjects. The analytical sample thus consisted of 47 whole blood samples from human subjects with definitive heart disease who were not experiencing an acute event at the time of sampling.

Details of the metabolite quantification have been described previously (Trainor et al., 2017), but a brief overview is provided as follows. Plasma samples were prepared from whole blood and a recovery standard was added. Vigorous shaking was applied utilizing a GenoGrinder 2000 (Glen Mills, Metuchen, NJ) and methanol was added and to precipitate proteins. The extract containing small molecules was divided into five aliquots, four of which were analyzed using different platforms while the remaining aliquot was reserved. Two aliquots were analyzed by ultra-performance liquid chromatography-tandem mass spectrometry (UPLC-MS/MS) with negative and positive ion mode electrospray ionization (ESI). A third aliquot was also analyzed by UPLC-MS/MS with negative ion mode ESI and a method optimized for polar metabolite detection. The fourth aliquot was analyzed by gas chromatography-mass spectrometry (GC-MS). 1,032 chemical species were detected utilizing the multiple platforms in the analysis of the plasma samples. Of these, 590 compounds were identified by matching to authentic standards based on retention index, mass to charge ratio, and MS2 data; 73 were identified based on experimental data matched to curated databases; and 369 could not be confidently identified. As the original data dependent acquisition was conducted utilizing both acute event samples and stable heart disease samples, metabolites not detected in the stable heart disease samples were removed. Metabolites missing from greater than 70% of the samples or without compound identification were also removed, resulting in a final dataset with 522 metabolites across 47 samples. Minimum values were then imputed for the remaining metabolite relative abundances with missing data. As many of the metabolites exhibited approximately log-normal relative abundance distributions, metabolite abundances were log-transformed. Finally, the data was mean centered so that each metabolite’s relative abundance distribution was centered about zero.

To approximate the posterior distribution of *p*(**Ω**|**X**), a Markov Chain was generated of length 1,000 with a 250 iteration burn-in period. From each sample from the posterior distribution *p*(**Ω**|**X**), a partial correlation coefficient matrix was computed, yielding a simulated posterior distribution for the matrix of partial correlation coefficients.

## 3. Results

### 3.1. Simulation studies Ω

Results from the simulation studies given an autoregressive covariance structure are shown in Table 1 and Figures 2–3. In Figure 2 a graphical model representation of the underlying covariance structure is shown along with the graphical model representations of the sample covariance matrix and the concentration matrices estimated by the multiple techniques evaluated in this study. These figures were generated from a randomly sampled simulation study. GGM estimation by the Bayesian Graphical Lasso (BGL) and Adaptive BGL exhibited similar performance characteristics with respect to sensitivity, specificity, AUC, and F1 measure. The performance of the chemical structure informative adaptive BGL varied significantly based on the suitability of the informative prior distribution for shrinkage parameters. In the “good prior” case in which it is assumed that the structural similarity and data generating process were deterministically linked, the structure adaptive BGL demonstrated significantly higher sensitivity, specificity, AUC, and F1 measure than the other techniques. Conversely, in the “poor prior” case in which the relationship between the simulated structural similarity and the data generating process was masked by gaussian noise, average AUC and F1 measure were significantly lower for the structure adaptive BGL than the remaining techniques.

**Table 1:** Results of the AR(1) simulation studies. For each of the compared techniques, sensitivity, specificity, area under the receiver operating characteristic curve (AUC), and F1 measure are reported. Reported values represent the sample mean and standard deviation over the simulation study replicates.

| Technique | Sensitivity | Specificity | AUC | F1 Measure |
| --- | --- | --- | --- | --- |
| Bayesian Graphical Lasso (BGL) | 0.6792 ± 0.073 | 0.9896 ± 0.007 | 0.9740 ± 0.014 | 0.7786 ± 0.054 |
| Adaptive BGL | 0.6702 ± 0.079 | 0.9936 ± 0.006 | 0.9793 ± 0.014 | 0.7827 ± 0.062 |
| Chemical Structure Adaptive BGL (Good prior) | 0.9894 ± 0.019 | 0.9997 ± 0.001 | 1.000 ± 0.000 | 0.9938 ± 0.011 |
| Chemical Structure Adaptive BGL (Poor prior) | 0.8175 ± 0.068 | 0.8763 ± 0.033 | 0.9148 ± 0.036 | 0.6448 ± 0.066 |

**Figure 2:**
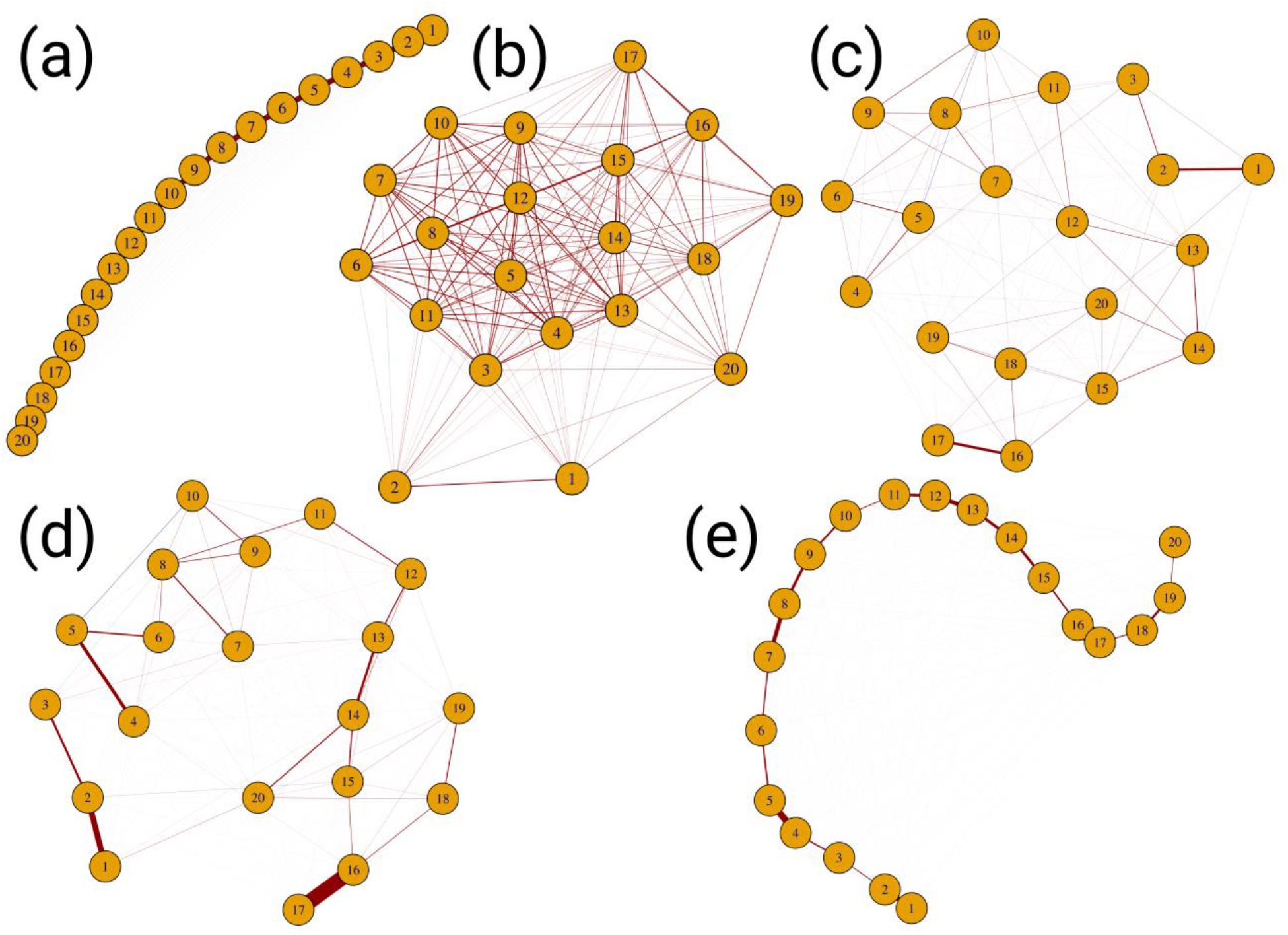
True and estimated concentration matrix graphs for a randomly selected AR(1) simulation study with *p* = 20 simulated random variates (metabolites) and a simulated sample size of *n* = 10. Graphs represent the: (a) true concentration structure given an AR(1) covariance structure with *p* = 0.95, (b) sample covariance matrix, (c) concentration matrix estimated by the non-adaptive BGL, (d) concentration matrix estimated by the adaptive BGL with non-informative priors, (e) concentration matrix estimated by the adaptive BGL with chemical structure informative priors.

**Figure 3:**
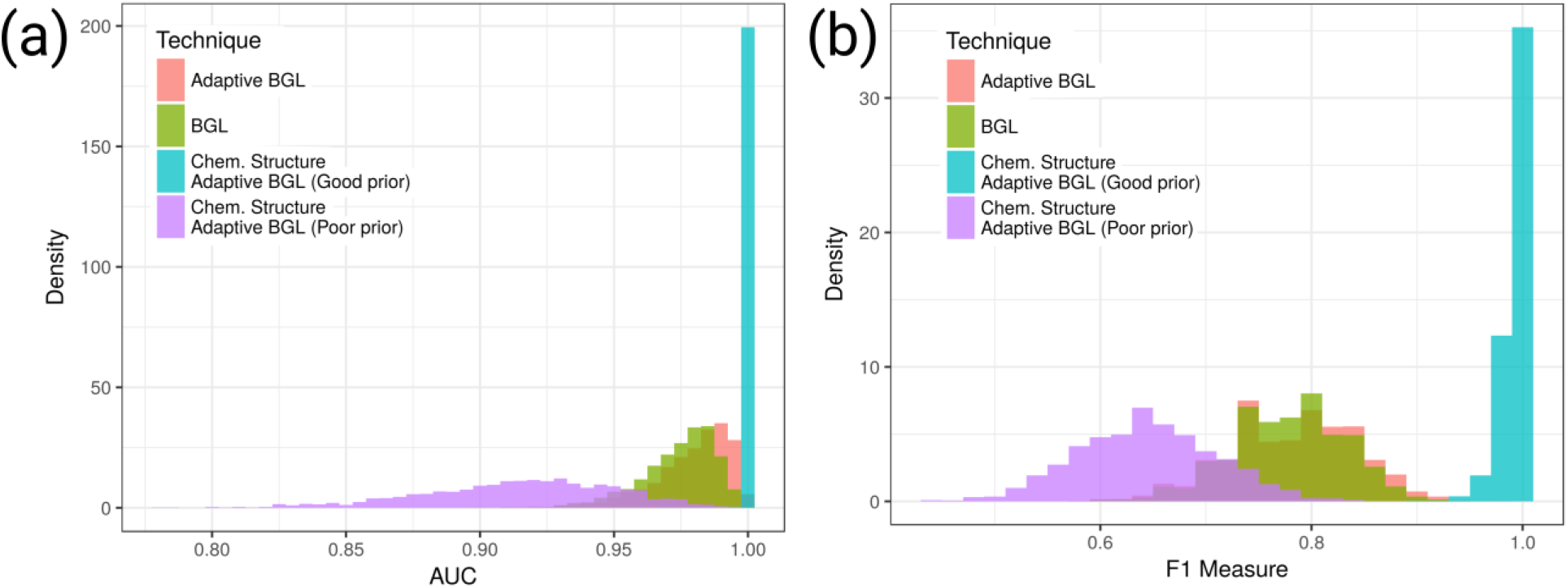
Results of the AR(1) simulation studies. The comparative performance of the techniques (as the ability to detect an edge, given an edge is truly present) is presented. Subfigure (a) shows a histogram of the observed area under the receiver operating characteristic curve (AUC) values, while (b) shows the F1 measure.

### 3.2. Plasma interactome for stable heart disease

A heatmap representation of the structural similarity between metabolites is shown in Figure 4. This heatmap was constructed using agglomerative hierarchical clustering using Ward’s method and squared distances (with distance computed as *d_ij_* = 1 – *s_ij_*, where *s_ij_* is the structural similarity between compounds). For illustrative purposes, the cluster containing cholate was retrieved from the root dendrogram by extracting the branch with height 0.6, as the structural-adaptive BGL subnetwork generated by cholate is considered later. Considering clusters generated by branches with low merge heights (high structural similarity), cholate was a member of a cluster with other closely related compounds such as deoxycholate, 3b-hydroxy-5-cholenoic acid, and glycocholate. Considering the more inclusive cluster generated by the branch at join height 3, other members included many intermediates in progestagen, androgen, glucocorticoid, and mineralocorticoid steroid metabolic pathways. These steroid hormone metabolites were all members of a cluster with similar within-cluster distances. Finally, at the same branch height that joined steroid hormone and cholate metabolites, a branch consisting of tocopherols cluster and squalene cluster was also joined.

**Figure 4:**
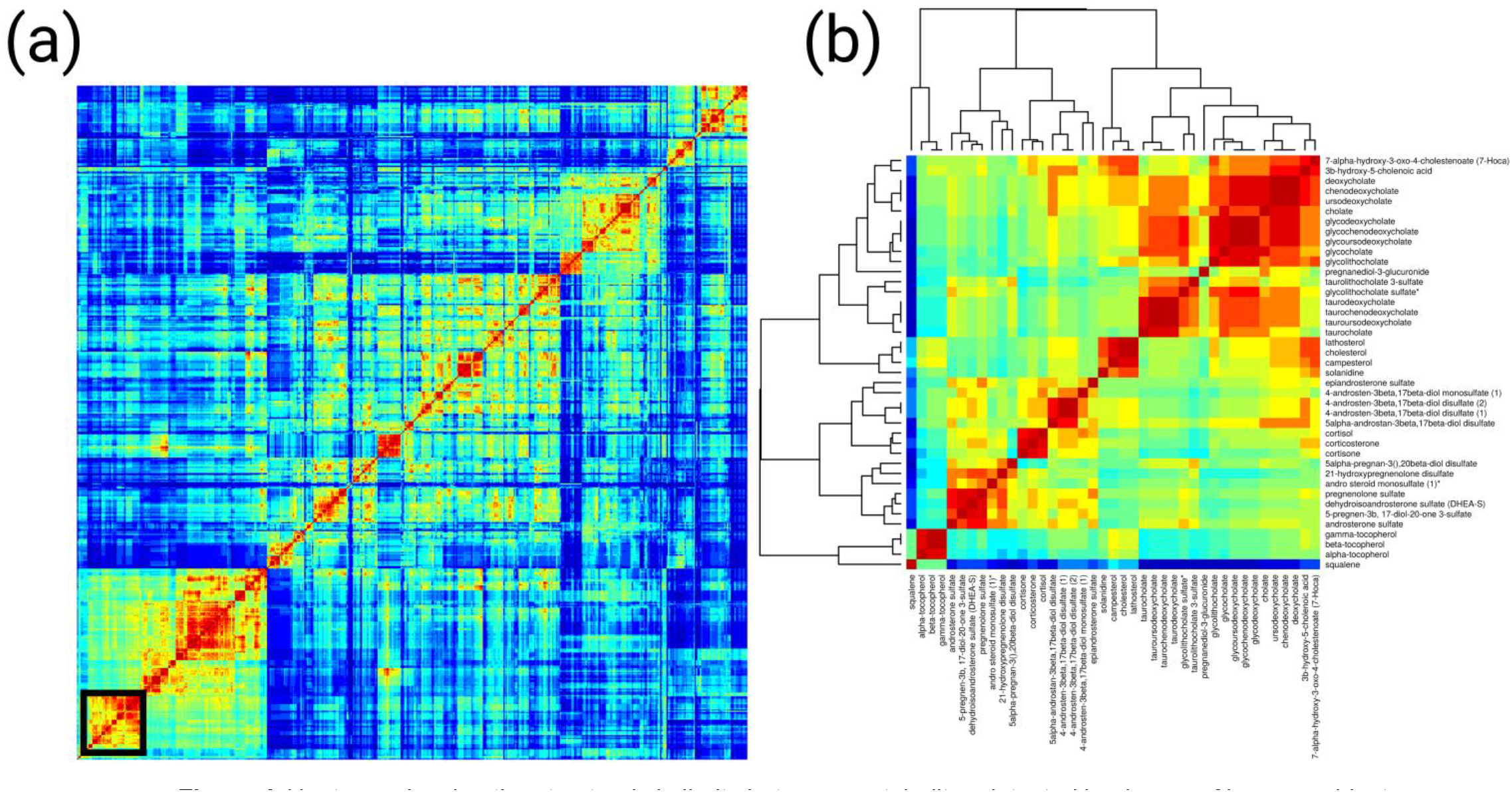
Heatmap showing the structural similarity between metabolites detected in plasma of human subjects presenting with heart disease. A local substructure coloring approach was utilized for quantifying similarity as the degree of overlap (presence and absence) of local structures based on atom types and bonds. Dendrograms were generated via agglomerative hierarchical clustering utilizing Ward’s minimum variance criterion and distances defined as 1 - similarity.

Elements of the MCMC sampling iterations are presented in Figure 5. Continuing with the working example of the metabolite cholate, Markov chains are presented for the estimation of the concentration parameters for the pairs (cholate, tyrosine), (cholate, cortisone), and (cholate, glychochenodeoxycholate). These metabolites are highlighted as exemplars of metabolites with relatively low, medium, and relatively high chemical structure similarity with cholate. The time series of the MCMC sampling for the shrinkage parameter, *λ_ij_*, demonstrated differences between the three pairs. Averaged across iterations, more shrinkage was applied with decreasing chemical similarity between the metabolite pairs. While the shrinkage parameter sample values for (cholate, cortisone) tended to be significantly smaller than the sample values for (cholate, cortisone), substantial overlap was observed in the posterior distribution of the concentration parameters for the same pairs.

**Figure 5:**
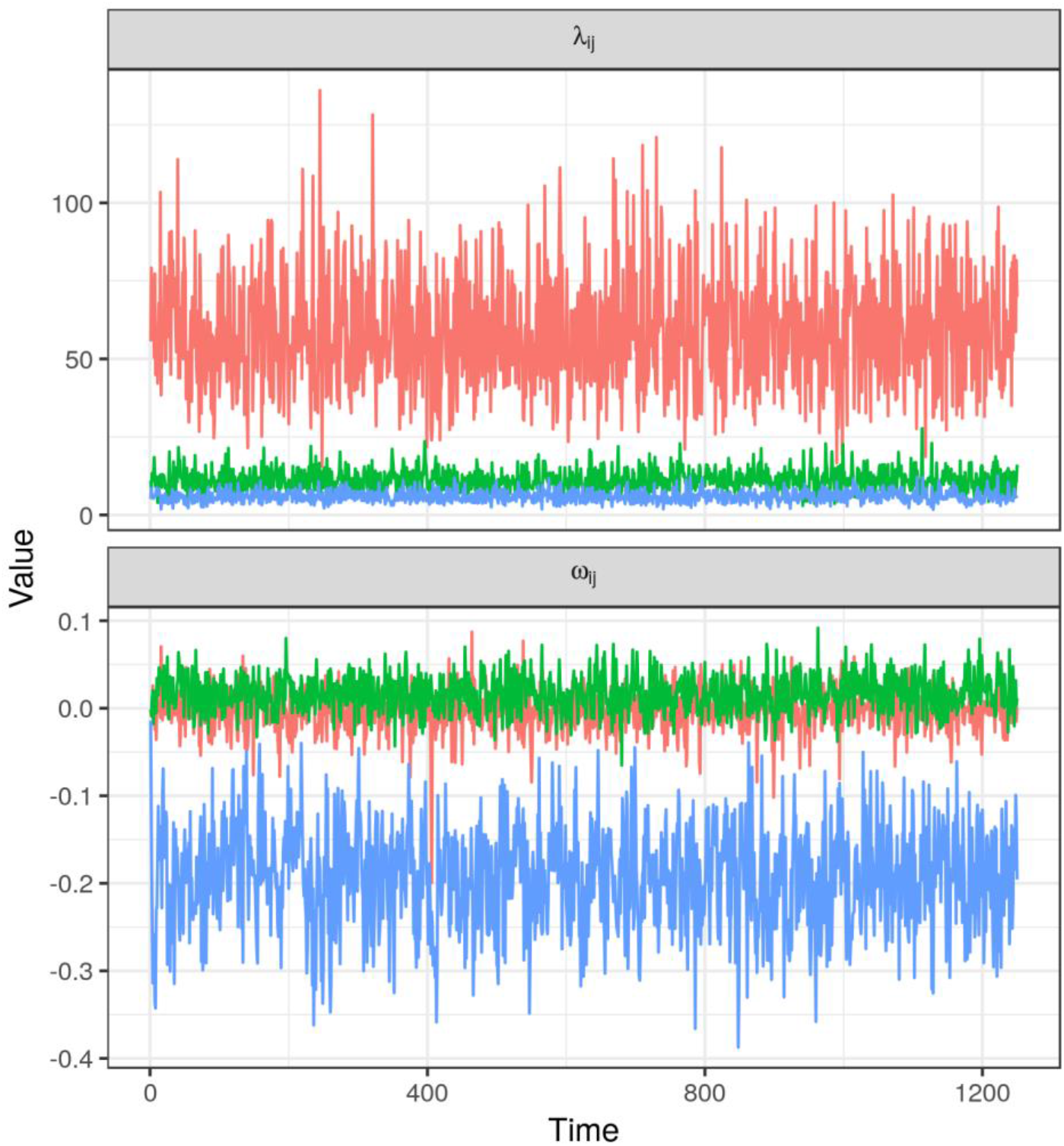
Time series plots for the MCMC sampler for the shrinkage parameter and *λ_ij_* for the concentration matrix entry **Ω**_*ij*_ for the following metabolite pairs: (cholate, tyrosine), (cholate, cortisone), (cholate, glycochenodeoxycholate).

From the simulated posterior distribution of **Ω**, the posterior mean *E*(**Ω**|**X**) was estimated after discarding burn in iterations. The posterior mean of the distribution of partial correlation coefficients was also computed. The resulting plasma metabolite interactome inferred by the structure-adaptive Bayesian Graphical Lasso is presented in Figure 6. This figure presents both the entire graph representing the posterior mean partial correlations as well as the subgraph generated by considering the neighbors of cholate. For ease of viewing, only edges for which |*ρ_ij_*| > 0.05 are plotted in the presentation of the full graph.

**Figure 6:**
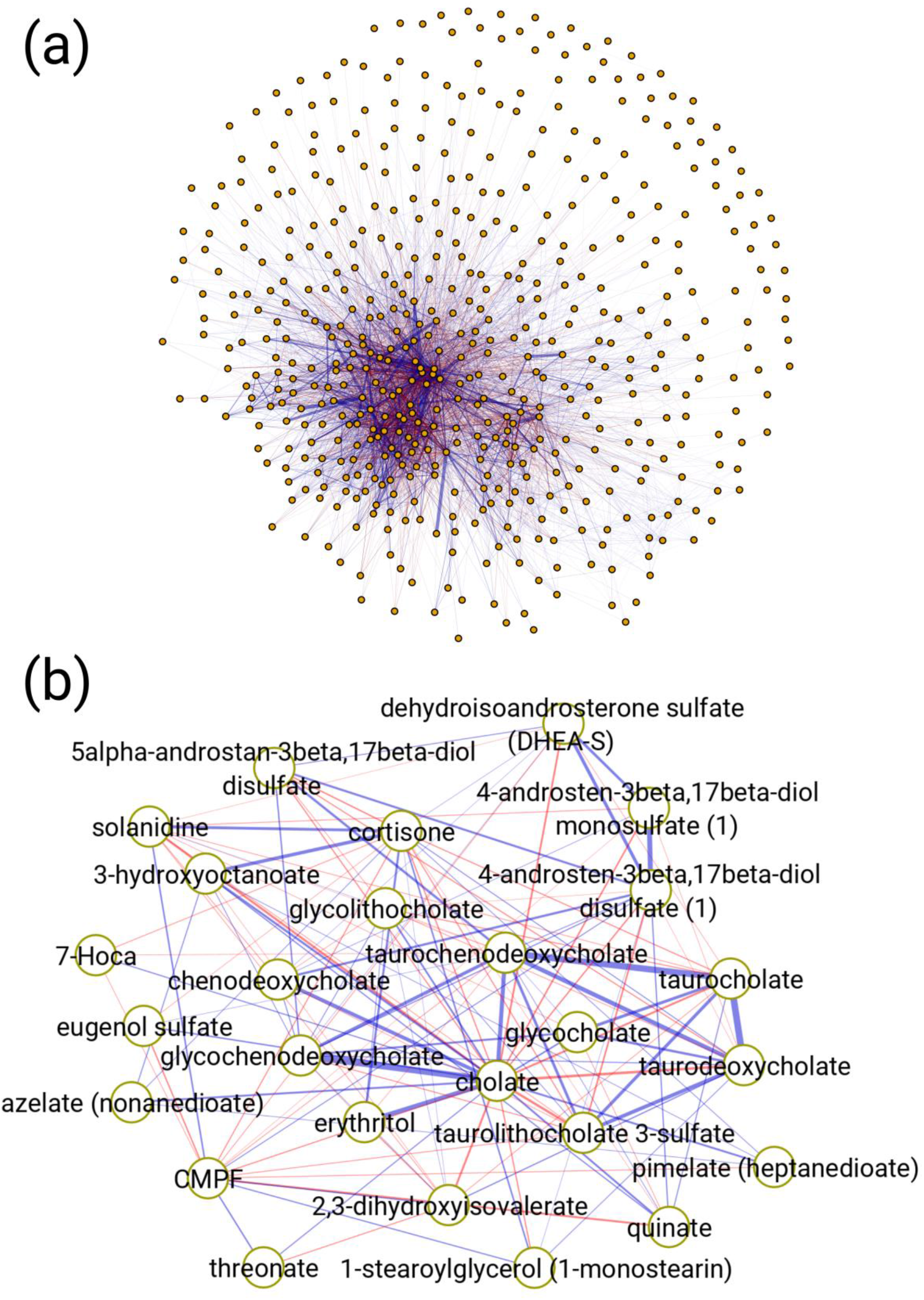
Graphical representation of the plasma metabolite interactome estimated by the chemical structure adaptive Bayesian Graphical Lasso (BGL) for stable heart disease. A simulated posterior distribution for the matrix of partial correlation coefficients was determined from the simulated posterior distribution of the concentration matrix **Ω**. Median values of each partial correlation coefficient were then determined and are represented as colored edges (negative values represented in red, positive values in blue). Subfigure (A) shows all metabolites in the interactome along with edges for which |*p*| > 0.05.

Positive partial correlation coefficients were observed between cholic acid the following other primary bile acids: glycocholic acid, chenodeoxycholic acid, and glycochenodeoxycholic acid. Negative partial correlation coefficients were observed between cholate and the following: taurocholic acid, taurodeoxycholic acid, and taurochenodeoxycholic acid. In addition, the bile acid 7-Hoca and the bile acid conjugate taurolithocholate 3-sulfate were first neighbors of cholic acid. Multiple conjugated androsterones were observed to be first neighbors of cholic acid as was the glucocorticoid cortisone and the steroidal alkaloid solanidine. Other metabolites that were first neighbors of cholic acid included: 3-Carboxy-4-methyl-5-propyl-2-furanpropionic acid (CMPF), eugenol sulfate, erythritol, 2,3-dihydroxyisolvalerate, threonate, quinate, pimelate, and azelate.

## 4. Discussion

Making inferences regarding how metabolic processes differ between phenotypes is the ultimate goal of most metabolomics and systems biology studies. Yet, unlike comparing the concentration or abundance of one metabolite across two or more phenotypes, for which simple statistical tests such as t-tests, Wilcoxon Rank-Sum tests, or multi-group analogues are readily available, a statistical framework for determining if and how metabolic processes differ between phenotypes remains elusive. Both strictly empirical methods (e.g. correlation analyses) and *a priori* knowledge based approaches (e.g. pathway enrichment analyses) suffer from substantial flaws. In terms of empirical methods, the analysis of correlations (such as by the Pearson, Spearman or biweight midcorrelation coefficient) reveals the marginal associations between metabolites; however, these methods do not uncover the relationship between a pair of metabolites conditional on the abundances of the remaining metabolites. Gaussian graphical models have been proposed previously in the context of metabolomics (Krumsiek et al., 2011) as an alternative, as GGMs can be utilized to determine the partial or conditional relationship between metabolites. In their work Krumsiek et al. (2011) show that GGM edges (or concentration matrix entries) estimated from the analysis of blood serum samples from a large human cohort correspond to known metabolic pathway interactions. Consistent with this, our simulation studies illustrate the advantage of analyzing metabolite-metabolite interactions using the partial correlation coefficients from a GGM as opposed to correlation networks. In the case of an autoregressive correlation structure as might be observed given a linear metabolic pathway, correlation networks exhibit extremely high connectivity, and consequently could not be utilized to elucidate the order of reactions. In contrast, we observe high sensitivity and specificity in detecting the true edges using a Bayesian Graphical Lasso estimated GGM. While the approach utilized by Krumsiek et al.(2011) was appropriate for the analysis of their data, it would not be possible to apply this approach in studies in which the sample size is smaller than the number of metabolites, as is common in many metabolomics studies. Frequentist regularization methods represent a class of solutions for ensuring that the concentration matrix, or equivalent GGM topology is estimable. In an implicit manner, frequentist regularization methods for estimating GGMs place a higher *a priori* probability on models with concentration matrix entries of smaller magnitude (H. Wang, 2012). However, this implicit prior cannot incorporate *a priori* knowledge as to whether some metabolites are more likely to be related than others.

In contrast to empirical methods, *a priori* knowledge based approaches such as pathway enrichment analyses consider the relationships between metabolites to be deterministically known which are then used to contextualize empirical results. Previous work (Dinesh Kumar Barupal & Fiehn, 2017; Dinesh K. Barupal et al., 2012) has highlighted that the coverage of metabolites detected in metabolomics studies in commonly utilized metabolic pathway and reaction databases may be extremely low. For example (Dinesh Kumar Barupal & Fiehn, 2017) observe that given 385 metabolites identified from the plasma of non-obese diabetic (NOD) mice, only 135 metabolites (or 35.1%) could be mapped to KEGG pathways (Kanehisa & Goto, 2000; Kanehisa, Sato, Kawashima, Furumichi, & Tanabe, 2016). To address this problem, Dinesh Kumar Barupal and Fiehn (2017) propose an alternative approach that utilizes both existing chemical ontological terms and chemical similarity between metabolites to develop coherent categories of metabolites for enrichment analyses. In the current work, we have sought a framework for balancing the benefits of empiricism with the benefits of *a priori* knowledge based approaches, while seeking to minimize the risks associated with both approaches. As opposed to considering metabolites as deterministically assigned to fixed pathways, our approach assumes that metabolites that are linked by biochemical reactions will exhibit overlap in local substructures. From this, our approach generates prior distributions for shrinkage parameters for the estimation of Gaussian graphical models. The posterior distribution of GGM parameters is thus proportional to the likelihood of the concentration matrix parameters (or the partial correlations between metabolites) times the prior probability of the concentration matrix parameters (which are linked to the structural similarity between metabolites). Similar to the non-informative BGL approach, this approach ensures that the concentration matrix is estimable via the Bayesian analog of regularization, however the regularization is applied given the prior belief that stronger associations are *a priori* more likely given structurally related compounds than unrelated compounds. While we find better justification for using structural similarity to generate prior probability distributions for shrinkage parameters in estimating a GGM, this approach would generalize to the use of priors from metabolic pathway maps. A previous work sought to estimate a GGM using 17 compounds quantified by NMR from 24 microglia cell culture samples using priors determined from KEGG (Peterson et al., 2013).

In addition to evaluation via simulation studies, we have applied the chemical structure adaptive BGL to generate a media-specific (blood plasma) metabolite interactome for stable heart disease. This model may serve as a reference model for comparing how the probabilistic interactions between metabolites in circulation change during acute disease events such as myocardial infarction or unstable angina. From this model, we have observed probabilistic interactions that are consistent with previous research in metabolism, as can be observed by focusing on the metabolite cholate. Bile acids are the major catabolic intermediate of cholesterol (Russell, 2003). Within mammals, the bile acid pool consists of primary bile acids such as cholic acid and chenodeoxycholic acid which are synthesized from cholesterol by enzymes expressed in hepatocytes, as well as secondary bile acids that are synthesized from primary bile acids by bacteria in the gut (García-Cañaveras, Donato, Castell, & Lahoz, 2012; Hofmann, Hagey, & Krasowski, 2010; Russell, 2003). In addition to bile acids aiding in the digestion of nutrients in the gut, bile acids also act as signaling molecules that have been shown to regulate glucose and lipid metabolism (Ferrebee & Dawson, 2015; Khurana, Raufman, & Pallone, 2011). Given the substantial proportion of cholesterol that is converted to bile acids leading to elimination, bile acid metabolism is linked to atherosclerosis (Meissner et al., 2013). In addition bile acids as signaling molecules affect cardiac (Desai et al., 2017; Rainer et al., 2013) and circulatory physiology (Khurana et al., 2011) via direct effects such as taurodeoxycholic acid mediated vasodilation (Khurana, Yamada, Wess, Kennedy, & Raufman, 2005). With respect to the current work, we observed relatively strong partial correlation between cholic acid and other primary bile acids. Additionally, partial correlations were observed between cholic acid and steroid hormones that share cholesterol as a common precursor. Given the importance of bile acids in cholesterol metabolism, atherosclerosis, cardiac physiology and circulatory physiology, a reference model of the probabilistic interactions of bile acids in circulation can help elucidate how acute disease events impact bile acid metabolism.

As with any Bayesian approach, the choice of prior probability distribution has a direct influence on the posterior distribution of model parameters (Gelman, 2014). In the current work, we have utilized informative priors that are linked to chemical structure similarity. This represents a potential limitation of the current work. Over the course of the simulation studies, we observed, unsurprisingly, that by introducing random noise into the simulated structural similarity the performance of the chemical structure adaptive BGL deteriorated. Further, the performance of the technique given “poor” prior information was, on average, worse than the performance of techniques such as the non-adaptive BGL that rely on non-informative priors. One element of the chemical structure adaptive BGL is worth noting in this context. In our proposed formulation, other monotonic functions for relating structural similarity to the Gamma scale parameter may be employed, as well as different shape and scale hyperparameters can be utilized. In this manner, the experimenter can diminish or strengthen the degree to which structural similarity impacts shrinkage. A second limitation of the current work is the choice of a multivariate Gaussian distribution for representing the joint distribution of metabolite abundances. While transformations in the stable heart disease data were applied over each metabolite to reduce the degree of departure from normality, the underlying intensity data is not normally distributed. Further, there are many cases in which approximate normality is not an achievable aim. A metabolite that is only present in some samples (e.g. acetaminophen metabolites that are present in some human subjects who have taken this medication, but not others) is one such case. Following a missing value imputation procedure, such a metabolite would exhibit a bimodal distribution that would not be well described by a Gaussian model.

